# Biofilm formation and maize root-colonization of seed-endophytic Bacilli isolated from native maize landraces

**DOI:** 10.1101/2024.01.02.573954

**Authors:** Gabriela Gastélum, Alejandra Ángeles, Guillermo Arellano-Wattenbarger, Yaxk’in Coronado, Eduardo Guevara, Jorge Rocha

**Author notes:** Address correspondence to Jorge Rocha.

## Abstract

Agricultural microbiology seeks to replace the use of agrochemicals with microbe-based products. Plant growth-promoting bacteria (PGPB) are often selected based on their functions *in vitro*, and then, their effect on plant development is tested. However, this approach neglects the study of their survival in soil, root-colonization, and the monitoring of beneficial functions in the rhizosphere. This could explain the recurrent lack of success in the transition from lab tests to field applications of natural isolates from novel habitats. In our recent studies, we found that native maize seeds from traditional agroecosystems carry endophytic bacterial communities that are dominated by members of the class Bacilli. As an approach to grasp their PGP potential, we developed protocols to test maize root-colonization of these natural isolates in 1) a short-term hydroponics assay *in vitro* and 2) a long-term assay in non-sterile soil pots. Our results show that *in vitro* biofilm formation was only partially associated to short-term colonization *in vitro*; furthermore, long-term root-colonization in soil pots was not correlated to the *in vitro* assays. This work highlights the necessity to incorporate root-colonization assays as part of the research strategies in the search for PGPB natural isolates from unexplored habitats, towards the generation of inoculants with increased success in the field.

## 1. Introduction

The use of plant-beneficial microbes in agriculture is a promising strategy to improve plant health and crop productivity while decreasing the use of agrochemicals. Bacterial strains from the genus *Bacillus* are abundant in soil, and are known for establishing diverse beneficial interactions with plants (Goswami, Thakker, and Dhandhukia 2016; Shafi, Tian, and Ji 2017). These interactions include plant growth-promotion (PGP) through the production of phytohormones, volatile organic compounds or by increasing the availability of nitrogen and phosphorus (Jiang et al. 2019; Nascimento et al. 2020; Sun et al. 2020). They also exhibit antagonistic activity towards plant pathogens by producing extracellular metabolites such as siderophores, antimicrobial compounds and hydrolytic enzymes; additionally, they can trigger induced systemic resistance in plants (Goswami et al. 2016; Mahapatra, Yadav, and Ramakrishna 2022; Miljaković, Marinković, and Balešević-Tubić 2020). For this reason, there is extensive literature exploring the beneficial effects of *Bacillus* isolates *in vitro* (Kalam, Basu, and Podile 2020; Rahmoune et al. 2017; Wahyudi et al. 2011), greenhouse (Akinrinlola et al. 2018; Masood, Zhao, and Shen 2020), and in field experiments (Kumar, Ahmad, and Singh 2022; Nguyen et al. 2019) using different plant species.

Research on plant-beneficial *Bacillus* normally focuses on their functions *in vitro*, and then the effect of microbial inoculation on plant growth is tested. However, the physical association of bacteria on the roots is often overlooked, despite the fact that bacterial establishment and persistence in the rhizosphere determine the development of plant-beneficial interactions (Parnell et al. 2016; Rilling et al. 2019). This is a major shortcoming in the field of agricultural microbiology as survival in soil and root-colonization are complex traits that depend on environmental conditions (Agler et al. 2016), edaphic factors (da Costa et al. 2020), root architecture (Birt et al. 2022), and interactions with the plant and native soil microbiota (Kurkjian, Akbari, and Momeni 2021; Wippel et al. 2021). As a result, the beneficial functions of bacterial inoculums observed *in vitro*, are often lost when applied in field (Bashan et al. 2014). In fact, root-colonization and mechanisms driving this association are frequently studied only after inoculants are commercialized (Kröber et al. 2014; Mendis et al. 2018; Xu et al. 2019).

Bacterial association with the host plant roots depend on biofilm formation (Knights et al. 2021); *i.e.*, the development of multicellular aggregates surrounded by an extracellular matrix that facilitates their adherence to surfaces (Branda et al. 2005). Despite the great diversity of plant-associated *Bacillus* spp. found in nature, research on biofilm formation and root-colonization mechanisms of this bacterial group is limited to only a few model bacteria, namely *B. subtilis* (reviewed in Blake, Christensen, and Kovács 2021). However, different strains may exhibit unique mechanisms to execute these processes, and the lack of low-cost and reliable methods to effectively track non-model natural bacterial isolates in the rhizosphere is a great limitation (Rilling et al. 2019). Therefore, new experimental strategies are needed to assess root-colonization in order to accelerate the discovery of natural isolates with PGP traits from unexplored habitats that display increased success in field.

In recent years, evidence has accumulated showing that many practices related to agricultural modernization have negative impacts on soil microbial diversity and plant-microbe interactions (Banerjee et al. 2019; Hartmann et al. 2015; Kavamura et al. 2018; Wang et al. 2020). For this reason, the search for isolates with novel plant-beneficial traits has focused on low-input agroecosystems (French et al. 2021). While exploring bacterial communities associated to native maize landraces from traditional *milpa* agroecosystems (Gutiérrez and Gómez 2011), we found that the seed-endophytic bacteriome of these plants is dominated by strains from the class Bacilli (*i.e*., *Bacillus* spp. and closely related sporulating bacteria) (Arellano-Wattenbarger et al. 2023; Gastélum et al. 2022). These bacteria are tightly associated with the plant, and we expect that they should be able to colonize the maize rhizosphere as a first step for their internalization into the endosphere and/or for establishing symbiotic interactions with the plant. Additionally, these strains may serve as models to explore novel mechanisms driving biofilm formation, root-colonization and beneficial functions.

Here we explore the relationship between *in vitro* biofilm formation and root-colonization of endophytic Bacilli isolated from native maize seeds, as a first approach to identify strains that display plant-beneficial functions. From our initial collection of isolates, we used 20 representative strains to assess biofilm formation *in vitro* and early root-colonization of maize *in vitro*; next, long-term root-colonization of 9 strains was tested in a more realistic soil pot assay. Although all these strains are tightly associated to maize seeds in nature, and many displayed biofilm formation and rapid colonization *in vitro*, only few were able to colonize the roots in our long-term assay in non-sterile soil. Noteworthy, our *in vitro* observations failed to predict strain performance in the soil pot assay. *Milpas* and other low-input agroecosystems may be ideal sources of a great diversity of beneficial microbes, but our results highlight the need of realistic experimental systems to study the beneficial functions of natural bacterial isolates as an initial step towards harnessing these genetic resources for sustainable agriculture.

## 2. Methods

### 2.1 Strains and growth conditions

Strains used in this study are shown in Table 1. *Bacillus* and *E. coli* strains were streaked from cryostocks in LB agar (10 g L^-1^ tryptone, 5 g L^-1^ yeast extract, 5 g L^-1^ NaCl and 15 g L^-1^ agar) and grown overnight at 30 °C or 37 °C, respectively. Inoculums were prepared by picking one colony into 5 mL of LB media and grown at 30 °C or 37 °C at 200 rpm for 16 h. Ampicillin (100 µg ml^-1^), erythromycin (5 µg ml^-1^), rifampicin (50 µg ml^-1^) or cycloheximide (10 µg ml^-1^) were added to the media when needed.

**Table 1.** Strains used in this study.

| Strain | Genetic background | Reference |
| --- | --- | --- |
| *Bacillus subtilis* NCIB 3610 | WT | (Branda *et al.*, 2001) |
| *Bacillus subtilis* PY79 | WT | (McLoon *et al.*, 2011) |
| *Escherichia coli* Top10 | WT |  |
| *Escherichia coli* | Carrying the plasmid pTMN387-*mKate2* | (Dragoš *et al.*, 2018) |

### 2.2 Maize seed collection

For the isolation of seed-endophytic spore-forming bacteria, maize seed samples of 8 different landraces (NH1, NH2, NH3, NH5, NH6, NH7, NV1 and NV2) from *milpas* in Hidalgo, México were previously collected (Gastélum et al. 2022). We also included two additional yellow maize landraces from different *milpas*: NH8, from Huitzizilingo, San Felipe Orizatlán, Hidalgo (21°10’25.504” N, 98°39’22.276” W), and NH11, from Atlaltipa Tecolotitla, Atlapexco, Hidalgo (21°00′02″ N, 98°22′09″ W). For early and long-term root-colonization assays, the hybrid variety YOMXA was used (distributed by MäHai seeds, Mexico).

### 2.3 Isolation of seed-endophytic spore-forming bacteria

We performed a selective isolation of spore-forming bacteria from either pulverized seeds, or roots of germinated seedlings. Seeds were surface-sterilized as described in previous work (Gastélum et al. 2022) and groups of 10 surface-sterilized seeds from each variety were pulverized in a sterile mortar or germinated in sterile conditions. For germination, seeds were placed on sterile paper towels soaked in 10 mL of sterile water inside a Petri dish. Seeds were incubated in the darkness for 3 days at 25 °C (Gastélum et al. 2022). Ten seedlings (or 10 pulverized seeds) were uses for obtaining bacterial suspensions in 10 mL of sterile PBS with a 30 s vortex agitation and soaking for 5 min. Each suspension was concentrated 10X and, in order to target spore-forming bacteria, samples were incubated at 80 °C for 20 min prior to plating 100 µL in LB. Inoculated plates were incubated at 30 °C for 5 days. Colonies with different morphologies were isolated and cryo-stocks for each strain were prepared using 15 % glycerol and stored at-80 °C.

### 2.4 Molecular identification and phylogenetic reconstruction

All the isolated strains were preliminarily identified by sequencing the V4 region of the 16S rRNA gene. This region was amplified using primers 515F (5’-GTG CCA GCM GCC GCG GTA A-3’) and 806R (5’-GGA CTA CHV GGG TWT CTA AT-3’) (Caporaso et al. 2011). Then, PCR products were purified using the PureLink Quick PCR Purification Kit (Invitrogen) and sequenced by the Sanger method (Unidad de Servicios Genómicos, Langebio-Cinvestav, Irapuato, Mexico). Genus-level identification was made using the Classifier from the Ribosomal Database Project (Cole et al. 2014) and a preliminary species-level identification was made using SeqMatch from the RDP. Species were assigned based on the type strain species showing the highest score with the query sequence. The phylogenetic tree was constructed using the sequenced region of all the strains, and the 16S rRNA gene of *Bacillus subtilis* NCIB3610 (ATCC accession number 6051) was included. All sequences were aligned using MAFFT v.7. (Katoh, Rozewicki, and Yamada 2017) with an automatically selected iterative refinement method. Since we used the complete sequence of the 16S rRNA gene of *B. subtilis* NCIB 3610, 553 and 771 positions were trimmed from the 5’ and 3’ ends of the alignment, respectively, to avoid gaps. The phylogeny was constructed in the online platform PhyML 3.0 using the Maximum Likelihood estimation and 1000 bootstraps. Smart model selection was used to determine the best substitution model (Guindon et al. 2010; Lefort, Longueville, and Gascuel 2017).

### 2.5 Biofilm formation assays

Biofilm formation was assessed through both qualitative and quantitative methods. For qualitative determination of biofilm formation, we used colony imaging. For this, 5 uL of stationary-phase culture (grown in LB broth at 30 °C for 16 h) were spotted on LB agar and on solid biofilm-inducing media Minimal Salts glutamate-glycerol (MSgg) (Branda et al. 2001). Biofilm formation was determined when colonies presented a complex architecture on MSgg compared to colonies on LB after 72 h of incubation at 30 °C. For the qualitative evaluation, we used the crystal violet assay with modifications (Merritt, Kadouri, and O’Toole 2011). Briefly, stationary-phase cultures were diluted 1:100 on MSgg liquid media. Then, 100 μL of the diluted culture were transferred into sterile 96-well microtiter plate (not tissue culture treated) in six replicates. The inoculated microtiter plate was incubated at 30 °C and biofilm formation was assessed after 72 h. After removing planktonic cells and washing the plate (all plate washes were performed as described in Merritt et al. 2011, biofilms in each well were stained with 250 μL of 0.1% (w/v) crystal violet for 10 min at room temperature. After staining, the crystal violet was removed, plates were washed two more times and the stained biofilms were solubilized using 250 μL of 30 % (v/v) glacial acetic acid per well with a 15 min incubation. For quantification, 125 μL of the solubilized stained biofilm were transferred to a clear flat-bottom microtiter plate, and optical density at 600 nm (OD_600_) was measured using a plate reader (Varioskan Lux, Thermo Scientific). Wells with non-inoculated MSgg were used as control.

### 2.6 Early root-colonization assay *in vitro*

We assessed early root-colonization of seed-endophytic Bacilli from native maize landraces in a hydroponic system. The development of this protocol required the standardization of disinfection of the seedling root to reduce microbial load that interfere in the quantification of inoculated bacteria, time of exposure to bacteria, and washes for bacterial detachment from the seedling roots for CFU counts. First, maize seeds were surface-sterilized and germinated in sterile conditions as described previously. Seedlings with a root length of 2 to 3.5 cm were disinfected with a solution of 0.0625% sodium hypochlorite (NaOCl) to reduce the effect of seed-endophytic microbes on the quantification of root-colonization (*i.e.,* microbes liberated during germination and that remained on the roots). For disinfection, seedlings were placed in a sterile breaker and submerged in the NaOCl solution for 10 min with constant manual agitation. Then, seedlings were washed twice with sterile water for 10 min. To eliminate any remaining NaOCl, seedling roots were individually rinsed for 15 s with sterile water. Seedlings were transferred to a sterile test tube (10 mm x 100 mm) with 1.8 mL of Minimal Salts Nitrogen Glycerol media (MSNg) (Beauregard et al. 2013) inoculated with 0.2 mL of stationary-phase bacterial culture at OD_600_= 0.2, ensuring a complete coverage of the root and avoiding contact with the seed. Seedlings were incubated at 30 °C and 90 rpm, and early root-colonization was quantified after 8 h of incubation. To remove non-attached cells, seedling-roots were washed for 30 s with sterile PBS under the stream of a washing bottle. Next, to detach root-colonizing cells for CFU quantification, seedlings were transferred individually to a 50 mL tube with 1 mL of PBS with Tween 20 (0.05% v/v) and seedlings were washed with a 30 s vortex agitation. 10-fold serial dilutions of the resulting suspension (containing the root-attached cells) were plated on LB agar. Inoculated plates were incubated at 30 °C overnight. To normalize data, the root length of each seedling was measured and root-colonization was expressed in CFU/cm of root.

### 2.7 Rifampicin resistant variants

To successfully detect and quantify long-term root-colonization of seed-endophytic Bacilli strains from soil, we generated rifampicin resistant variants of nine strains. First, 40 mL of stationary-phase cultures of each strain were centrifuged at 8000 rpm and 4 °C for 10 min, and bacterial pellets were suspended in 2 mL of fresh LB media. For strains BsPY79, NME_32, NME_117 and NME_239, bacterial pellets were suspended in 0.7 mL of fresh LB media. Then, 500 μL of each bacterial suspension were plated on LB media with rifampicin. Plates were air dried and incubated in darkness for 48 h at 30 °C. After incubation, 3 to 10 rifampicin resistant (Rif^r^) colonies of each strain were isolated. Variants were checked for colony morphology (LB agar), colony architecture (MSgg agar), and growth rate (LB liquid media) in comparison with the corresponding parental strains. Selected strains were grown on LB liquid media and preserved with 15% glycerol at-80 °C.

### 2.8 Long-term root colonization in soil pots

Soil used on this assay was sampled from a monoculture system in San Salvador, Hidalgo (20°18’54.5“N 99°00’33.2”W). For quantification of long-term root-colonization, first, seeds were sowed in germination trays and watered daily. After 15-18 days, plants were transferred to pots with 300 g of soil and inoculated with rifampicin resistant (Rif^r^) Bacilli strains. For inoculation, Rif^r^ strains were streaked from cryostocks in nutrient agar plates (NA, 8 g L^-1^ Difco™ Nutrient Broth, BD, and 15 g L^-1^ agar), and incubated for 48 h at 30 °C. Then, 2 to 5 colonies were streaked as a lawn on NA plates and incubated 3 days at 30 °C. After incubation, bacterial biomass was recovered and suspended in 5 mL of sterile Phosphate Buffer Saline (PBS). Then, spores and vegetative cells were counted using a Neubauer chamber, and 1×10^7^ Rif^r^ CFUs were suspended in 70 mL of PBS and inoculated in each pot. After inoculation, plants were maintained in a growth chamber with 14 h of light and 10 h of dark at an average temperature of 25 °C for 15 days. Plants were watered every 3 days with 70 mL of tap water. After 15 days, plants were removed from their pots and soil excess on the roots was manually removed. Then, the following morphometric values were assessed: root and shoot fresh weight, stem length and total root length. Total root length was measured using the RhizoVision Explorer software (Seethepalli and York 2020). To quantify root-colonization, roots were separated from shoots using a sterile scalpel and rhizosphere and rhizoplane were sampled. To sample the rhizosphere, complete roots were transferred to a 50 mL tube and washed with 25 mL of sterile PBS. Tubes were slowly inverted 10 times, following 2 vortex pulses of 5 s each and 10 s of vigorous manual agitation. The resulting suspension was designated as the rhizosphere fraction. Then, roots were rinsed with PBS to discard any remaining non-attached cells. To sample the rhizoplane, roots were transferred to another tube and washed with 25 mL of PBS supplemented with Tween20 (0.05% v/v) to detach strongly associated bacteria. Tubes were vortexed with 2 pulses of 10 s each followed by 10 s of vigorous manual agitation. The resulting suspension was designated as the rhizoplane fraction. Finally, total CFUs on the rhizosphere and rhizoplane were calculated by plating 10-fold serial dilutions of the corresponding samples on LB media with rifampicin. In addition to rifampicin, chyclohexamide was used to inhibit fungal growth. Data was normalized using the total root-length of each plant assessed and values were expressed as CFU/cm of root. Not-inoculated plants were used as control. At least 8 plants per treatment were evaluated.

## 3. Results

### 3.1 Collection of seed-endophytic Bacilli from native maize landraces

In this work, we studied seed-endophytic strains from native maize landraces that are released during germination and remain on the surface of seedling roots. We isolated 32 spore-forming strains from 10 maize landraces and preliminarily identified them through partial 16S rRNA sequencing (Fig. 1). All isolated strains were classified into the phylum Bacillota (formerly Firmicutes). At the genus level, 15 strains were classified as *Bacillus* sp., 11 as *Paenibacillus* sp., 3 as *Peribacillus* sp. and 3 as *Domibacillus* sp. (Fig. 1), all belonging to the class Bacilli. For the following experiments, we selected a subset of 20 strains based on their phylogeny and the maize landrace of origin (Fig. 1), aiming at selecting a representative and diverse group.

**Figure 1.**
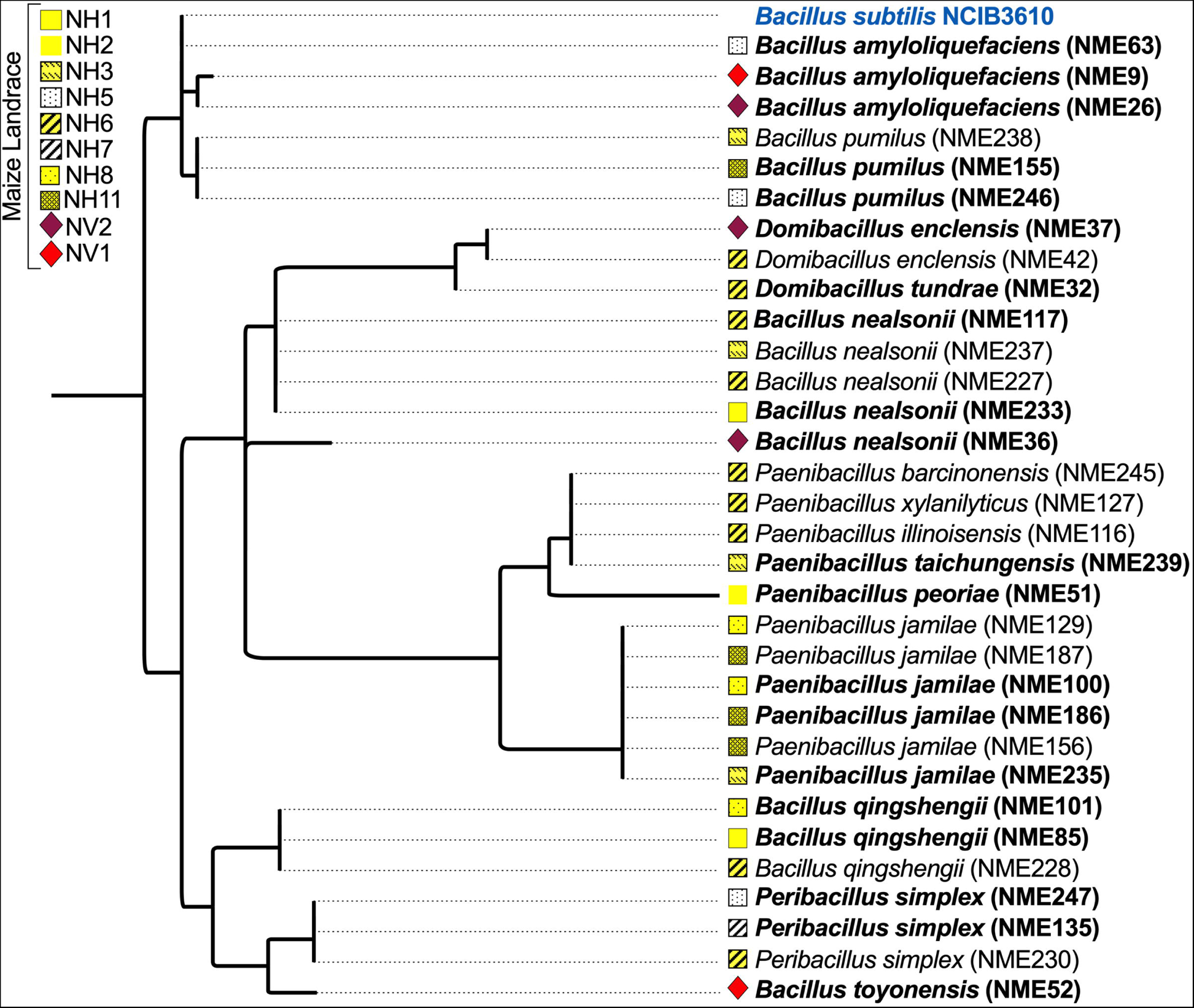
Phylogenetic tree of seed-endophytic Bacilli isolated from native maize landraces. *Bacillus subtilis* NCIB3610 is highlighted in blue and strains selected for early root-colonization assays are shown in bold. Strain codes are shown in parentheses.

### 3.2 *In vitro* biofilm formation of seed-endophytic strains from maize landraces

Previous studies show that genes involved in biofilm formation of *B. subtilis in vitro* are also necessary for root-colonization (Beauregard et al. 2013; Dragoš et al. 2018). Hence, biofilm formation is an important trait for the PGP capabilities of this model bacterium. As a first approach to address root-colonization of Bacilli strains from native maize, we evaluated biofilm formation *in vitro*. Qualitatively, we evaluated the colony architecture of strains in biofilm inducing agar media (Msgg) (Branda et al. 2001), and in nutrient-rich agar media (LB). For a quantitative evaluation, we performed the crystal violet assay (Merritt et al. 2011). For both assays, the biofilm forming strain *B. subtilis* NCIB3610 (Bs3610) and the domesticated laboratory strain *B. subtilis* PY79 (BsPY79) were used as reference (McLoon et al. 2011).

Bs3610 displayed colony architecture when grown in Msgg media, while BsPY79 did not (Fig. 2a). In the case of the seed-endophytic Bacilli, only 3 strains showed complex colony architecture in Msgg media: NME_63, NME_9 and NME_26, all of which were identified as *B. amyloliquefaciens* (Fig. 2a). In the crystal violet assay, Bs3610 displayed higher biofilm formation compared to BsPY79 (mean OD600 of 1.59 and 0.1, respectively) (Fig. 2b). Seed-endophytic strains NME_63, NME_9 and NME_26 (which displayed complex colony architecture, Fig. 2a) presented a high biofilm formation of 8.17, 9.01 and 5.22 mean OD600, respectively (Fig. 2b). Additionally, strains NME_155, NME_246, NME_37 and NME_51, showed an intermediate biofilm formation (mean OD600 of 0.9, 0.63, 0.73 and 0.35, respectively). The remaining 13 strains from native maize did not exhibit biofilm formation in this assay. Of note, all strains presented planktonic growth in in these static cultures (not shown). These results show that colony architecture in Msgg media and biofilm formation in the crystal violet assay were correlated only for Bs3610 and three seed-endophytic maize strains (NME_63, NME_9 and NME_26). For strains NME_155, NME_246, NME_37 and NME_51, biofilm formation in the crystal violet assay was not associated to the presence of a complex colony architecture in agar media.

**Figure 2.**
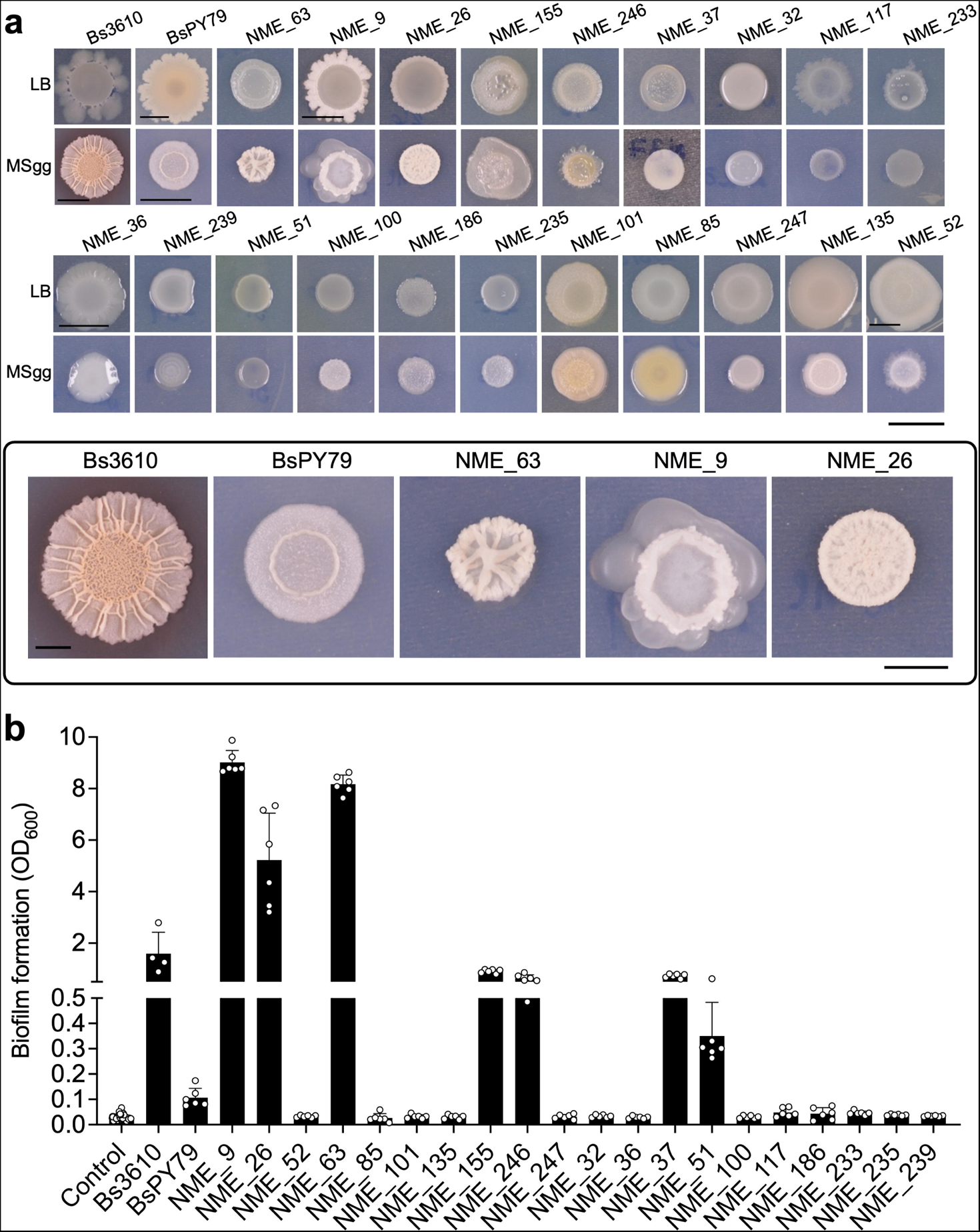
*In vitro* biofilm formation. a, qualitative assessment of colony architecture. Scales: 1 cm. Box: zoomed-in pictures of control strains and seed-endophytic strains showing complex colony architecture on MSgg media; scale: 0.5 cm. **b,** quantification of biofilm formation using the crystal violet assay. Not inoculated MSgg was used as control. Error bars indicate SD.

### 3.3 Early colonization of maize seedling roots

In studies that test plant growth-promotion by natural bacterial isolates, root-colonization is normally not tested (Rilling et al. 2019) mainly due to the lack of cost-effective and practical protocols for non-model bacteria-host experimental systems. We developed a protocol to quantify early root-colonization of maize in a hydroponic system (Fig. 3a). Briefly, this protocol consists of: 1) germinating seeds in sterile conditions, 2) disinfecting seedlings to reduce native microbial load, 3) exposing seedlings to Bacilli isolates, 4) seedling washes to obtain cells that strongly attach to the roots, and 5) CFU counts (Fig. 3a; see materials and methods).

**Figure 3.**
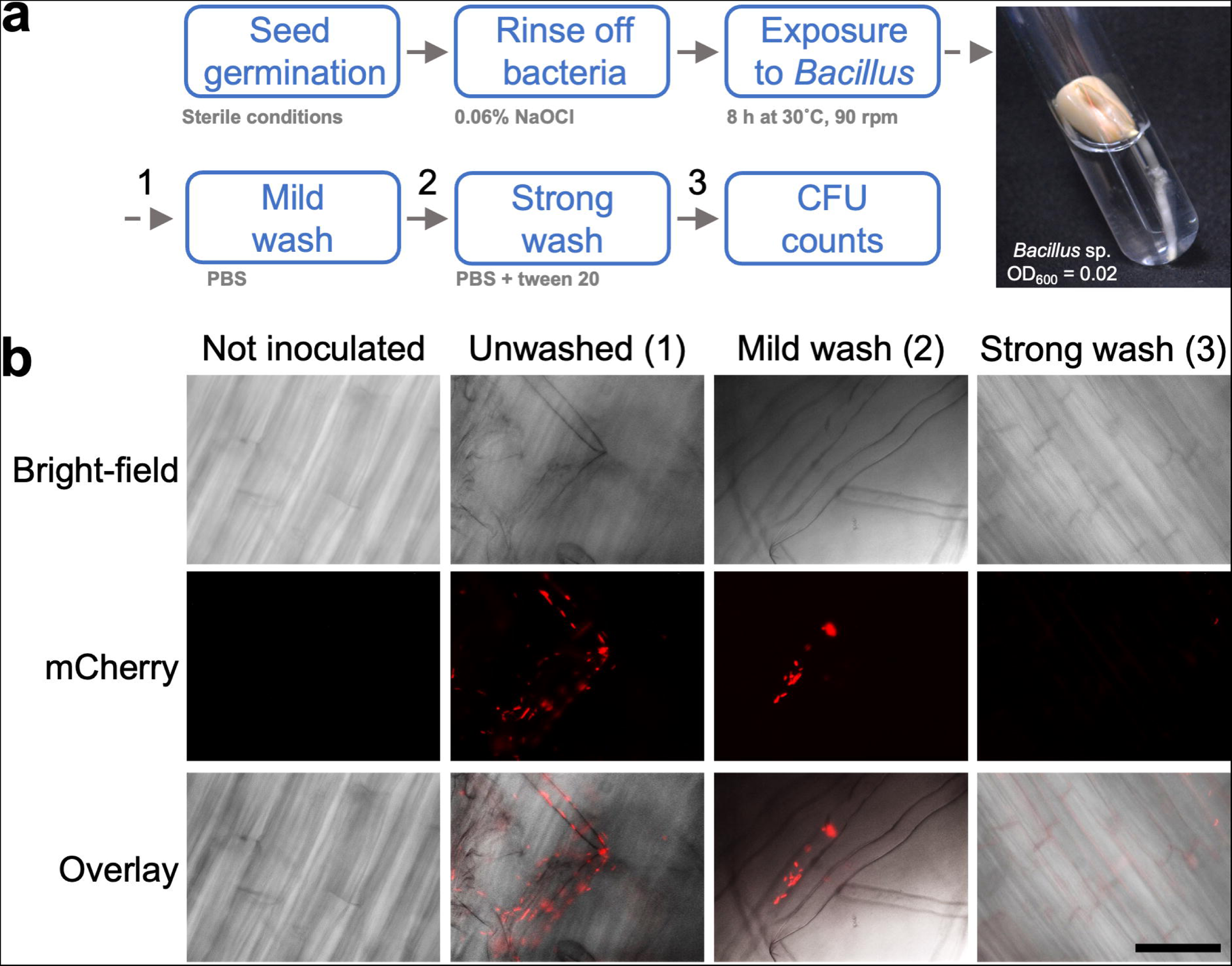
Method for the quantification of early root-colonization in maize seedlings. a, Flow chart representation of the method developed to quantify early root-colonization in maize seedlings. Numbers above arrows indicate steps in which roots were visualized with fluorescence microscopy. **b,** Seedling-roots inoculated with *E. coli* pTMN387-*mKate2* viewed with fluorescence microscopy after 8 h of incubation. Treatments 1, 2 and 3 correspond to steps indicated in part a. Objective: 20X, Scale: 50 μm.

In preliminary assays, we found that *E. coli* Top10 – which was used as a reference strain from an unrelated habitat – exhibited root-colonization in this system; therefore, we used a fluorescent strain *E. coli*-mKate to directly assess colonization under fluorescence microscopy in the following stages: 1) directly after bacterial exposure, 2) after a mild wash to discard non-attached cells and 3) after a strong wash to detach root-colonizing cells. After 8 h of bacterial exposure, fluorescent signal was abundant on and near the roots (Fig. 3b). After the mild wash, cells were only detected in patches directly on the root surface; finally, after the strong wash, cells were not detected (Fig. 3b). These results support that our protocol is useful to quantify cells that strongly attach to the roots after 8 h of exposure.

We quantified early root-colonization of the 20 Bacilli strains from native maize (Fig. 1), as well as the reference strains Bs3610, BsPY79 and *E. coli* Top10. As shown in figure 4, all tested strains were detected above the counts in not-inoculated seedlings (1.39×10^2^ CFU/cm) which corresponded to remaining seed-endophytic bacteria. Strains Bs3610, BsPY79 and *E. coli* Top10 showed an early root-colonization of 9.36×10^5^, 8.94×10^4^ and 2.34×10^6^ CFU/cm, respectively. Seed-endophytic Bacilli strains were able to colonize roots in a range from 4.78×10^3^ to 8.69×10^6^ CFU/cm (Fig. 4). Strains NME_36, NME_186, NME_246 and NME_155 displayed the highest early root-colonization (8.69×10^6^, 8.42×10^6^, 5.78×10^6^ and 5.25×10^6^ CFU/cm respectively), while strains NME_233 and NME_85 exhibited the lowest early root-colonization (2.2×10^4^ and 4.7×10^3^ CFU/cm, respectively). In this experiment, seedlings were exposed to bacterial cultures adjusted to an Optical Density of 600 nm (OD600) of 0.02; although this OD corresponded to varying cell concentrations of each strain, early root-colonization was not influenced by the cell density in the bacterial suspension used for seedling inoculation (Supplementary Fig. S1). These results show that Bacilli strains from native maize display early root-colonization in this *in vitro* assay, but this capability is highly variable between isolates.

**Figure 4.**
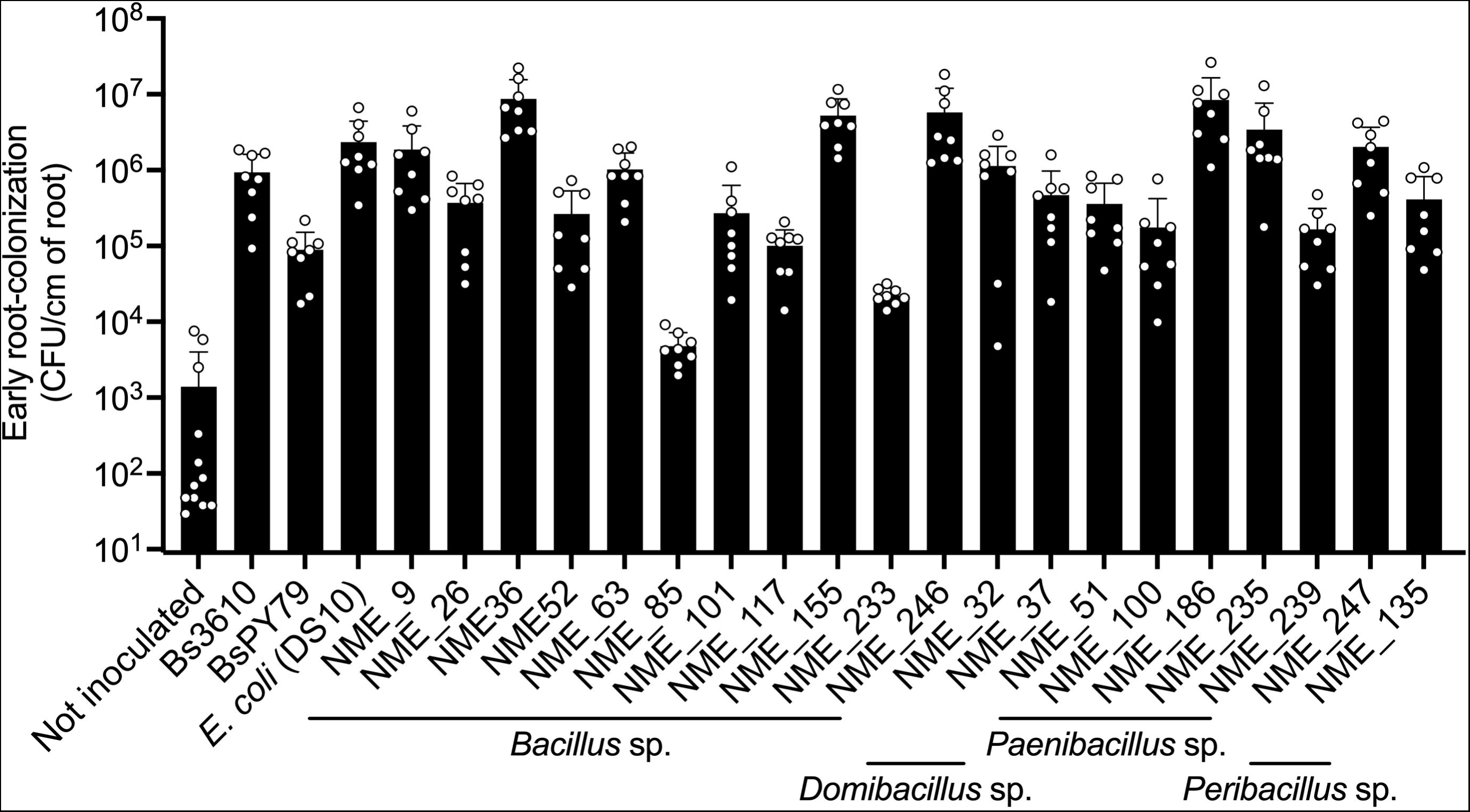
**Quantification of early root-colonization by seed-endophytic Bacilli**. Not inoculated seedlings were used as control. Dots represent measurements of individual plants. Error bars indicate SD.

Next, we addressed the relationship between *in vitro* biofilm formation and root-colonization, which is extensively acknowledged for the model bacteria *B. subtilis* (Beauregard et al. 2013; Blake et al. 2021; Dragoš et al. 2018). We plotted the corresponding values of each strain in a coordinates graph and found that this relationship is partially maintained since all biofilm forming strains were able to colonize roots in the hydroponics assay (Fig. 5); similarly, some non-biofilm forming strains exhibited deficient early root-colonization. However, we also found a group of strains that did not form biofilms *in vitro* but were capable of colonizing roots, some with values higher than the reference strain Bs3610 (*e.g.,* NME_247, NME_235, NME_36 and NME_186, Fig. 5). This suggests that some Bacilli strains from our collection may colonize roots through mechanisms that differ from those described in model laboratory strains. Also, that the crystal violet assay in Msgg media, which was developed for a model *B. subtilis* strain (Branda et al. 2001), is not adequate for quantification of *in vitro* biofilm formation of diverse natural isolates. The relationships found between early root-colonization (ERC) and *in vitro* biofilm formation (BF) allowed us to classify the 20 strains into three groups: group I (BF –, ERC –), group II (BF –, ERC +) and group III (BF +, ERC +) (Fig. 5) for subsequent experiments.

**Figure 5.**
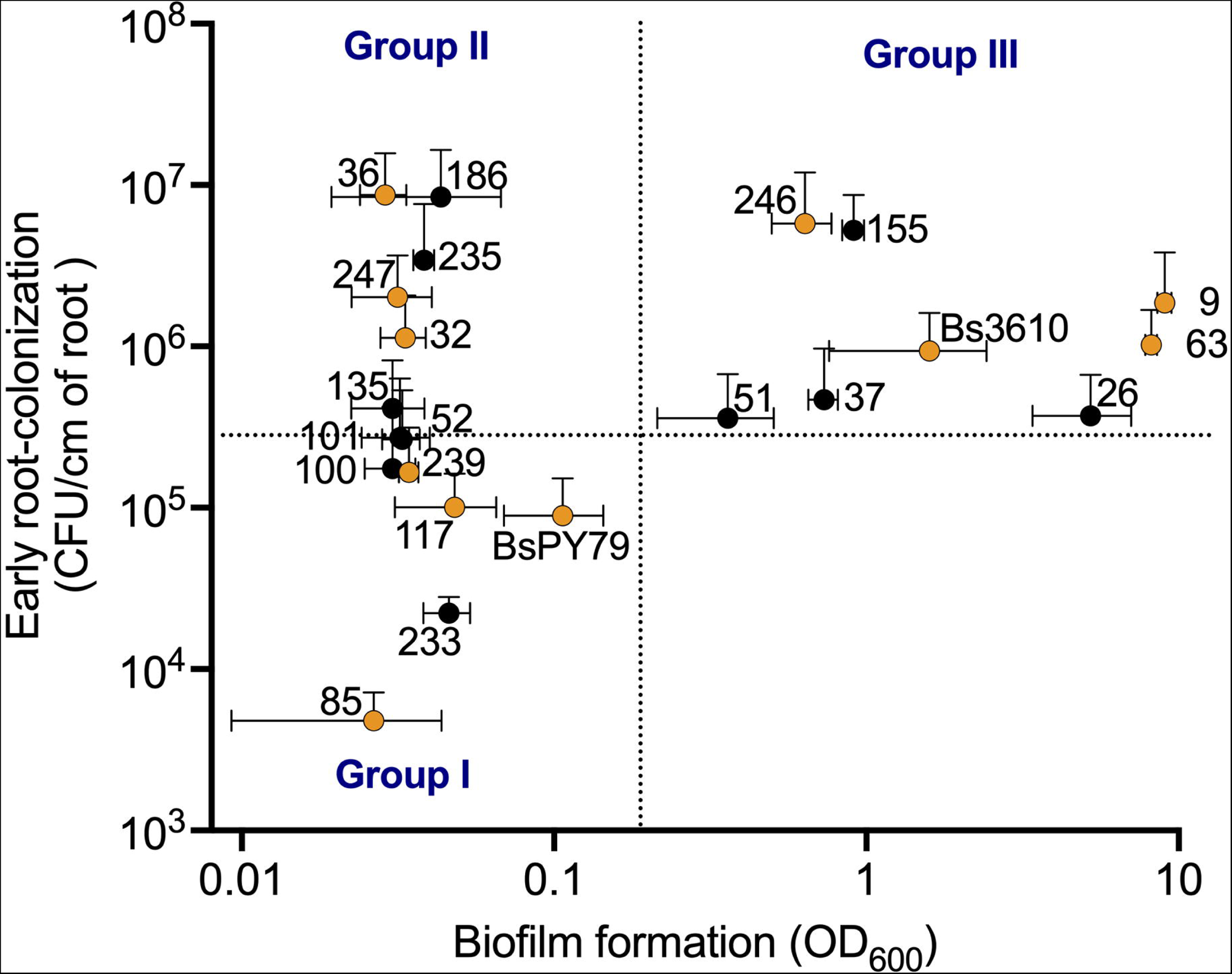
Correlation between biofilm formation and early root-colonization of seed-endophytic Bacilli from native maize landraces. Numbers represent the code of each NME strain. Three groups of strains were identified according to their capacity to form biofilms and early colonize roots. The groups enclosed with dotted lines. Biofilm Formation (BF), Early Root-Colonization (ERC). Group I: BF –, ERC –; Group II: BF –, ERC +; Group III: BF +, ERC +. Strains selected for long-term root-colonization assays are indicated with yellow dots. Error bars indicate SD.

### 3.4 Long-term colonization of maize plants in soil

In nature, microbial root-colonization is influenced by complex interactions with local soil microbiota and the host plant in a dynamic fashion during plant growth (Edwards et al. 2018; de Souza et al. 2016) which are not considered in our *in vitro* assay (Fig. 4). In order to address colonization of the Bacilli strains in a more realistic setting, we developed an experimental system to evaluate long-term root-colonization using maize plants grown in non-sterile soil. For this, we selected nine Bacilli isolates (three from each group previously identified; Fig. 5) as well as the two reference strains Bs3610 and BsPY79; from these strains, we generated spontaneous rifampicin resistant variants (Rif^r^) to facilitate their detection from the rhizosphere. First, we verified that this antibiotic prevented bacterial growth from non-inoculated soil (Supplementary Fig. S2). Then, we corroborated that CFU counts were not affected by the soil matrix (*e.g.* due to cell aggregation or adsorption on soil particles) using samples inoculated with known concentrations of a Rif^r^ strain (Supplementary Fig. S3).

We developed a method for quantifying the Rif^r^ strains from the rhizosphere and rhizoplane of mature maize plants (Fig. 6a). In this assay, 2-week-old maize plants were inoculated with individual strains at ∼3×10^4^ CFU/g of soil and colonization was addressed 2 weeks later. CFUs were not detected in non-inoculated plants (Fig. 6b). Strains Bs3610 and BsPY79 were able to colonize both the rhizosphere (1.8×10^3^ and 2.4×10^3^ CFU/cm, respectively), and the rhizoplane (2.78×10^2^ and 5.96×10^1^ CFU/cm, respectively) (Fig. 6b). Unexpectedly, we found that long-term root-colonization of native maize strains was not consistent with the phenotypes of *in vitro* biofilm formation and early root-colonization (Fig. 5). From group I (BF –, ERC –), strains NME_117 and NME_239 were not detected on neither the rhizosphere nor rhizoplane fraction (Fig. 6b); however, strain NME_85 was found in both fractions (3.89×10^2^ CFU/cm in rhizosphere and 1.35×10^1^ CFU/cm in rhizoplane) (Fig. 6b). From group II (BF –, ERC +), strains NME_36 and NME_247 were capable of colonizing the rhizosphere (2.46×10^2^ and 1.04×10^2^ CFU/cm, respectively) and rhizoplane (8.92×10^1^ and 4.8 CFU/cm, respectively); while strain NME_32 was not detected (Fig. 6b). From group III (BF +, ERC +), strains NME_9 and NME_63 colonized the rhizosphere only in half of the tested plants, with a mean root-colonization of 8.9 and 2.5 CFU/cm, respectively. These strains were not detected in the rhizoplane. Strain NME_246 from group III colonized both fractions with a mean root-colonization of 6.94×10^2^ and 2.73 ×10^2^ CFU/cm in the rhizosphere and rhizoplane, respectively.

**Figure 6.**
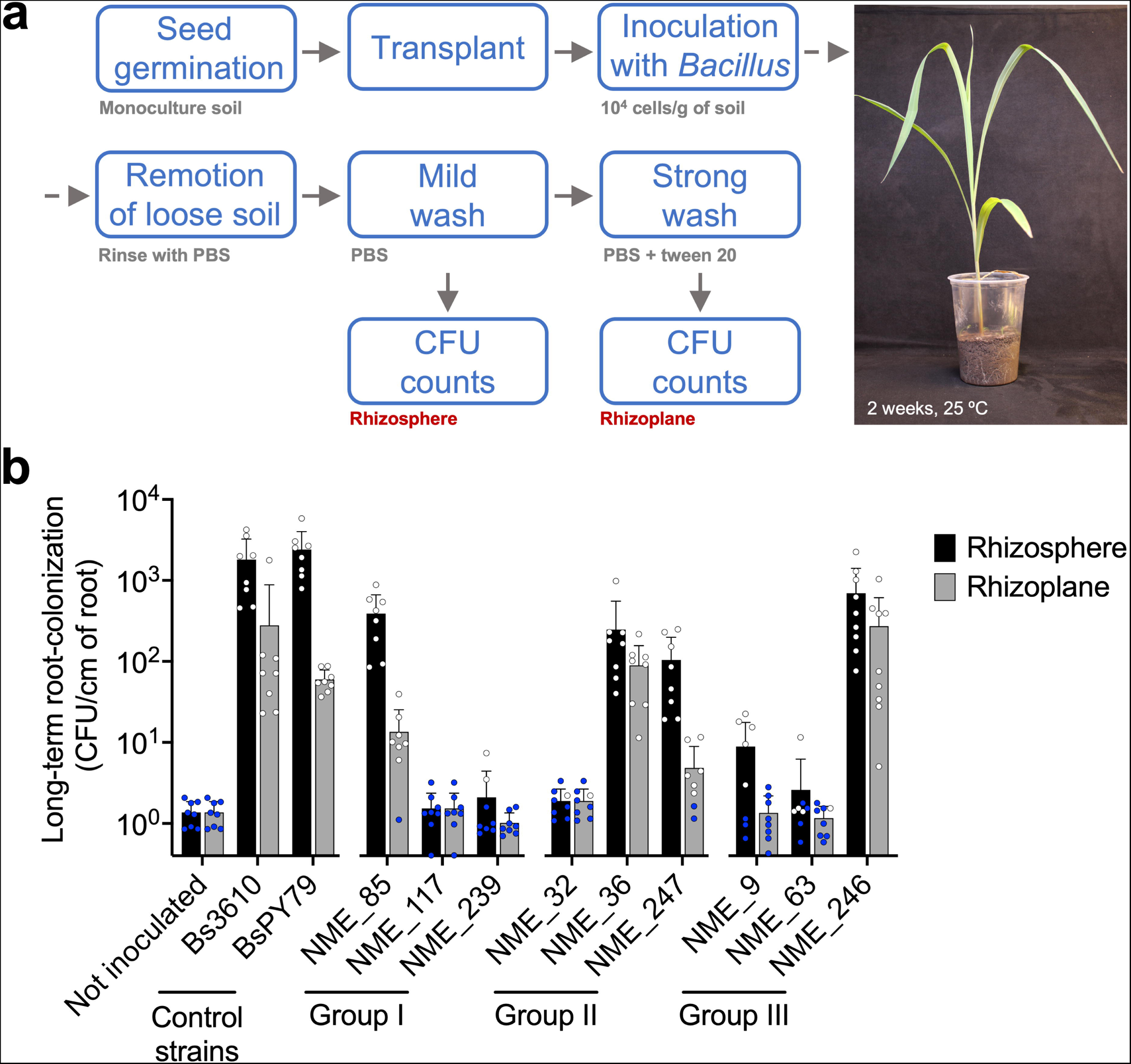
Long-term root-colonization by seed-endophytic Bacilli. a, Flow chart representation of the method developed to quantify long-term root-colonization. **b,** Quantification of long-term root-colonization. Dots represent independent samples. Blue dots represent samples with undetected CFUs. When CFUs were undetectable, values were considered as the detection limit of the assay for each case (1 CFU/root length). Bacterial groups appear as designated in figure 5: Group I (BF –, ERC –); group II (BF –, ERC +); group III (BF +, ERC +). Error bars indicate SD.

Survival in soil is a key determinant for bacterial inoculates to initiate the root-colonization process in presence of root exudates (Kaminsky et al. 2019). To discard that root-colonization results were solely influenced by the capacity of these strains to survive in bulk soil, we aimed at detecting the inoculated bacteria in bulk soil from six experimental pots where three colonizing and three non-colonizing strains were inoculated. All the non-colonizing strains tested (NME_9, NME_117 and NME_239) were successfully detected in bulk soil (Supplementary Fig. S4). Only one colonizing strain (NME_36) was not detected in bulk soil. We also verified that long-term root-colonization was not correlated to sporulation efficiency of the strains, which was measured upon inoculation (Supplementary Fig. S5). These results indicate that long-term root-colonization was not exclusively influenced by their survival in bulk soil due to higher sporulation; instead, their detection in the rhizosphere or rhizoplane may depend on other more complex phenomena such as their ability to reach the roots and establish interactions with the plant and the native soil microbiota.

All Bacilli tested displayed unique patterns in the phenotypes of *in vitro* biofilm formation, early and long-term root-colonization (Table 2). These patterns were unexpected as neither *in vitro* biofilm formation nor early root-colonization were associated with long-term root-colonization of these strains (Supplementary Fig. S6). We only found a positive correlation between rhizosphere and rhizoplane colonization in the soil pot assay (Supplementary Fig. S7), which was expected since these data were generated from the same experimental plants.

**Table 2.**
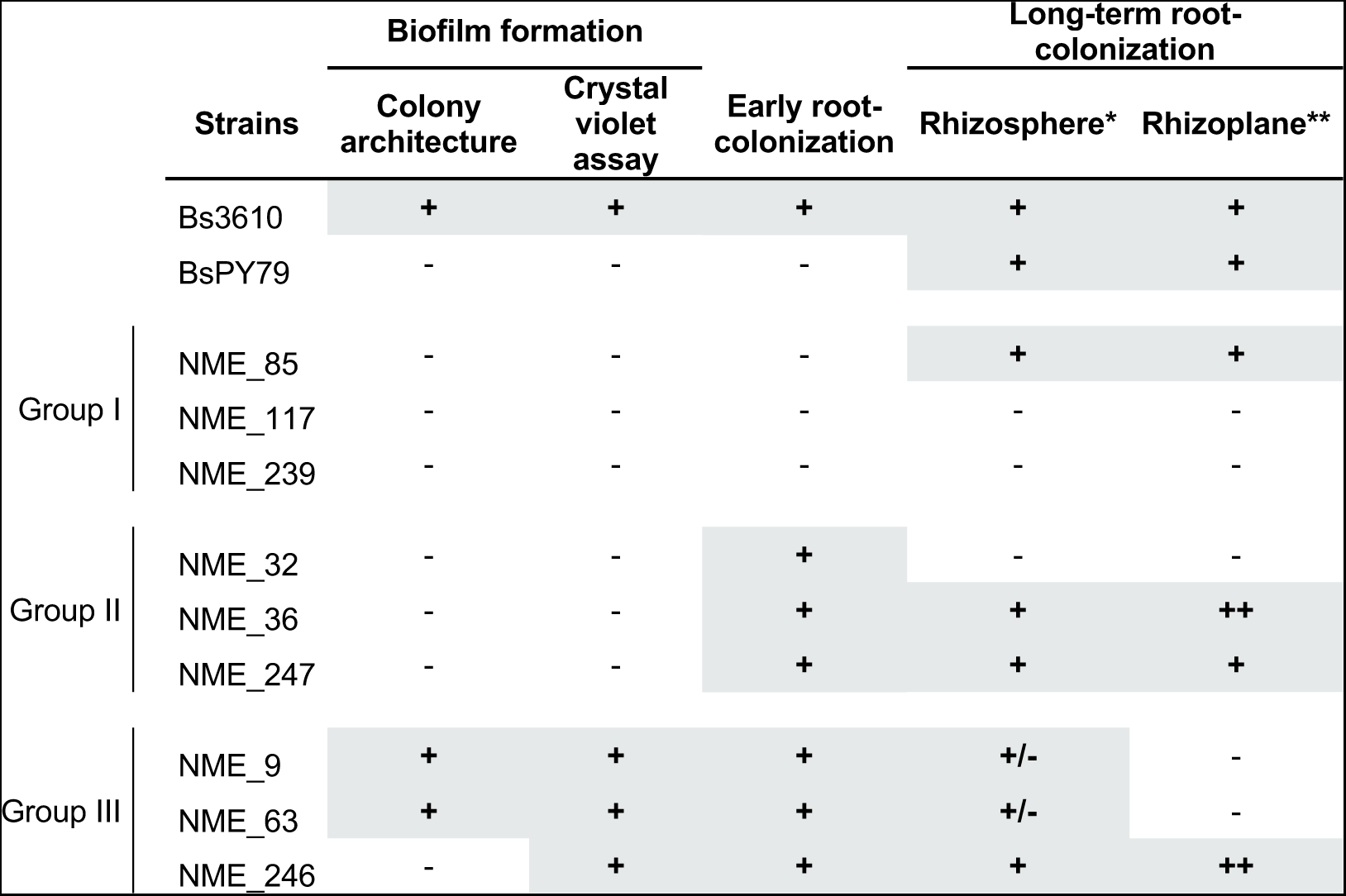
Summary of biofilm formation and root-colonization of seed-endophytic strains from maize landraces.

Finally, even though these assays were not intended for testing plant growth-promoting (PGP) traits of the native maize Bacilli, we collected morphometric measurements from inoculated plants. Although some Bacilli from native maize exhibited a successful long-term root-colonization, inoculation with these strains did not exhibit plant growth-promotion when compared to non-inoculated plants (Supplementary Fig. S8). Further experiments are being directed to test PGP traits of strains that are capable of colonizing roots, using longer assays with more experimental replicates and higher inoculation doses.

## 4. Discussion

Here we studied the relationship between *in vitro* biofilm formation, early root-colonization *in vitro,* and long-term root-colonization in soil, using a collection of natural isolates of seed-endophytic Bacilli strains from native maize landraces. Root-colonization is determinant for the establishment of beneficial plant-microbe interactions; likewise, understanding the environmental, ecological and molecular factors driving this complex function is key for harnessing microbial PGP traits (O’Callaghan, Ballard, and Wright 2022; Rilling et al. 2019). It is then conflicting that research on mechanisms underlying root-colonization, and its association to PGP traits, is normally limited to few bacteria-host model systems (Blake et al. 2021; Monteiro et al. 2012). By developing simple and cost-effective experimental systems to quantify root-colonization, we found that biofilm formation *in vitro* is only partially associated to short-term root-colonization in our natural isolates; furthermore, these two *in vitro* assays were not useful to predict rhizosphere and rhizoplane colonization in our long-term assay in soil pots. Therefore, incorporating the evaluation of root-colonization in realistic experimental settings during the selection of novel bacterial isolates as plant growth-promoting bacteria (PGPB), should lead to the discovery of strains with better performance in natural settings.

We proposed that seed-endophytic Bacilli from native maize could serve as model strains to test root-colonization as an early step to identify PGP strains, since these are tightly plant-associated bacterial communities. All 20 strains attached to the roots in our short-term hydroponics assay *in vitro*. This assay may resemble the germination stage since we used seedlings in a nearly axenic setup. Hence, this result supports that these bacteria are the first colonizers of the roots, and in turn they could contribute to the assembly of microbial communities in the rhizosphere (Figueiredo dos Santos et al. 2021; Gastélum et al. 2022; Shao et al. 2021). In contrast, only four out of nine strains were able to successfully colonize the rhizosphere and rhizoplane in the long-term assay in soil pots; in addition, two strains were detected only in the rhizosphere. Root-colonization in nature is a highly complex process, involving multiple steps such as survival in soil, spore germination, chemotaxis and motility towards the root, metabolization of root-exudates, interactions with the native microbiota, and biofilm formation (revieviewed in Knights et al., 2021). Genome sequencing could provide further insights into the physiological processes and molecular mechanisms associated to root-colonization of these natural isolates, which could explain poor colonization in the soil assay. Additionally, it has been shown that some collective functions such as motility and biofilm formation emerge in the context of particular bacteria-bacteria interactions (McCully et al. 2019; Yannarell et al. 2019), hence, the use of consortia or synthetic communities could be useful to study colonization of these Bacilli by considering bacterial interactions.

There is a vast and continuously growing literature testing PGP traits of a wide variety of bacterial strains (see Supplementary Table S1 for a non-comprehensive list including studies in maize, wheat and rice); nevertheless, the transition from lab tests to field applications is often unsuccessful (De Bruin et al. 2010; Kaminsky et al. 2019; O’Callaghan et al. 2022). As a result, after decades of research on PGPB we are still far from substituting chemical inputs with microbial-based products in agriculture; we propose that this is partly due to the poor understanding of the performance of these strains in natural settings, and specifically because root-colonization is only tested in a minority of studies (Rilling et al. 2019). Additionally, in the few cases where inoculated strains are tracked, experimental protocols often require high-priced equipment and/or present technical limitations when studying non-model bacteria. These include immunological techniques such as ELISA and immunofluorescence (Hansen et al. 1997; Yegorenkova et al. 2010) the use of reporter genes including the β‒galactosidase, β‒glucoranidase, bacterial luciferase and green fluorescent protein genes (Compant et al. 2005; Kragelund, Hosbond, and Nybroe 1997; Solanki and Garg 2014), and nucleic acid-based methods like FISH (Watt et al. 2006). More recently, microscopy (Hulse 2018; Liu et al. 2021), microfluidic devices (Aufrecht et al. 2018; Massalha et al. 2017; O’Neal, Vo, and Alexandre 2020) and genomic approaches (Kröber et al. 2014) have been used. Here we developed simple culture-dependent protocols based on direct CFU counts to assess root-colonization of non-model Bacilli, which will be useful for the study of bacterial colonization in other non-model bacterial-host experimental systems.

The diversity and functions of plant-associated microbes is affected by modern agricultural practices (Banerjee et al. 2019; Hartmann et al. 2015; Kavamura et al. 2018; Wang et al. 2020); for this reason, research on plant-microbe interactions in low-input agroecosystems is promising in the search for PGPB. *Milpas* are low-input traditional agroecosystems found in Mesoamerica where native maize landraces are cultivated in polyculture with other crops and wild plants (Gutiérrez and Gómez 2011). The soil and plant-associated microbiome from *milpas* is highly complex and diverse (Aguirre-Von-Wobeser et al. 2018; Van Deynze et al. 2018; Rebollar et al. 2017); specifically, seed-endophytic bacterial communities of native maize are more abundant and diverse when compared to hybrid varieties, and include strains that could contribute to biotic and abiotic stress alleviation (Arellano-Wattenbarger et al. 2023; Gastélum et al. 2022). From this fraction of the bacteriome, the class Bacilli appears particularly important, since they are a dominant taxon in these communities; however, their contribution to plant fitness remains elusive. After the soil pot assay, plants did not show significant changes in their development due to inoculation. Since this assay was aimed at evaluating colonization, we used 10^4^ cells/g of soil to inoculate pots, which is 100 to 1000-fold lower that the amount normally used in inoculation experiments aimed at evaluating PGP traits (Efthimiadou et al. 2020; Kálmán et al. 2023). Additionally, the experimental period was insufficient to address the effect of inoculation in plant developmental stages that impact productivity such as flowering, pollination and grain filling (Aslam et al. 2015; Nafziger 2016). Future experiments addressing PGP potential will focus on strains NME_85, NME_36, NME_247 and NME246, which display root-colonization, using increased biological replicates, higher inoculation doses and longer experimental periods, as well as applying stressful biotic and abiotic conditions (Hernández-Canseco et al. 2022; Jin et al. 2021; Omae and Tsuda 2022).

## 5. Conclusions

Meeting the nutritional needs of the growing global population requires a significant increase in crop productivity (Tian et al. 2021), and excessive chemical and water inputs related with modern agriculture cause several human health and environmental damage. In the last decades, agricultural microbiology has facilitated successful applications for more sustainable crops, including nitrogen fixation (Dent and Cocking 2017), plant growth (Dobbelaere et al. 2001) and pest biocontrol (Chattopadhyay, Bhatnagar, and Bhatnagar 2004; Raddadi et al. 2007). However, successful microbial inoculants in the field requires a deep understanding of the ecological and molecular mechanisms driving the establishment and persistence of introduced microbes in the rhizosphere. Our results highlight the need to study potential PGPB in more realistic experimental setups, in order identify strains with better performance in field. We propose that PGPB research is in an urgent need of a paradigm shift where studies include the evaluation of long-term tracking (detection) and monitoring (measuring the expression of beneficial functions) of natural isolates after plant inoculation. Likewise, more complex experimental model systems – such as microbial synthetic communities – should be developed more broadly to assess the effect of microbial interactions with the host and with the native microbiota on beneficial functions and root-colonization.

## Acknowledgements

Authors thank local producers from Hidalgo for providing native maize seeds for this study: José Antonio Hernández Hernández and Alfonso Lopez Bautista. Authors thank Paola Rodríguez for providing technical assistance. Gabriela Gastélum and Guillermo Arellano-Wattenbarger received a scholarship from CONAHCYT Mexico (761832 and 774643, respectively).

