## Supplementary for "Biofilm formation and maize root-colonization of seed-endophytic Bacilli isolated from native maize landraces"

[1] Unidad Regional Hidalgo. Centro de Investigación en Alimentación y Desarrollo A.C., San Agustín Tlaxiaca, Hidalgo, Mexico.

[2] Environmental Chemistry Department, Universidad Tecnológica de Querétaro. Santiago de Querétaro, Querétaro, México.

[3] Centro de Investigaciones Biológicas del Noroeste (CIBNOR). La Paz, Baja California Sur, México.

#### Supplementary Tables

**Table S1. Representative studies identifying natural bacterial isolates and the effect of their inoculation on the development of maize, wheat and rice plants.**

| Plant specie | Inoculated Bacteria | Variables measured | Optimal or stress condition | Reference |
|---|---|---|---|---|
| Maize | *Methylobacterium* sp. NMS14P | Growth measurements | Optimal | (Jirakkakul et al., 2023) |
| | *Bacillus subtilis*, *Bacillus megatherium,* and *Azotobacter chroococcum* | Growth and yield measurements | Optimal | (Efthimiadou et al., 2020) |
| | *Bacillus megaterium, Bacillus pumilus, Pseudomonas fluorescens, Pseudomonas putida* | Water use, water demand, germination, growth and yield measurement | Drought stress | (Kálmán et al., 2023) |
| | *Burkholderia phytofirmans, Enterobacter aerogenes, Pseudomonas fluorescens* | Gaseous exchange, biochemical, ionic, growth and yield measurements | Salt stress | (Afzal et al., 2023) |
| | *Pseudomonas putida, Pseudomonas fluorescens , Bacillus subtilis, Azospirillum lipoferum* | Biochemical, growth and yield measurements | Optimal | (Singh et al., 2023) |
| | *Bacillus spp., Azotobacter chroococcum, Priestia megaterium* | Biochemical, growth and yield measurements | Optimal | (Katsenios et al., 2022) |
| | *Azospirillum lipoferum* CRT1 | Germination, biochemical, growth and yield measurements | Optimal | (Rozier et al., 2017) |
| | *Pseudomonas fluorescens* | Biochemical, growth and yield measurements | Drought stress | (Zarei et al., 2020) |
| | *Azospirillum brasilense* Sp245, *Azospirillum brasilense* AbV5 + AbV6, *Herbaspirillum seropedica*e ZAE94. | Growth, yield, and nutritional measurements | Optimal | (Martins et al., 2018) |
| | *Azospirillum brasilense, Bacillus subtilis, Pseudomonas fluorescens* | Growth, yield, and nutritional measurements | Optimal | (Pereira et al., 2020) |
| Wheat | *Arthrobacter aurescens, Bacillus atrophaeus, Enterobacter asburiae, Pseudomonas fluorescens* | H+-PPase, HKT1, NHX7, CAT, and APX gene expression | Salt stress | (Safdarian et al., 2020) |
| | *Azospirillum brasilense* | Biochemical, growth and yield measurements | Optimal | (da Silva et al., 2022) |
| | *Bacillus wiedmannii* | Growth and yield measurements | Drought stress | (Karimzad et al., 2023) |
| | *Alcaligenes faecalis* LF.29, Bacillus sp LF.43 | Grain germination | Salt stress | (Almutairi et al., 2023) |
| | *Bacillus sp., Azospirillum lipoferum, Azospirillum brasilense* | Germination, gaseous exchange, biochemical, growth and yield measurements | Drought stress | (Akhtar et al., 2021) |
| | *Bacillus megaterium* TRQ8, *B. cabrialesii* TE3T, *B. paralicheniformis* TRQ65, *B. subtilis* TSO9 | Growth measurements | Optimal | (Robles Montoya et al., 2020; Rojas Padilla et al., 2020) |
| | *Azospirillum brasilense* | Growth and yield measurements | Optimal | (Diosnel et al., 2019) |
| | *Bacillus velezensis* | Growth and yield measurements | Optimal | (Nguyen et al., 2019) |
| | Serratia spp., Klebsiella spp. | Biochemical and growth measurements | Salt stress | (Acuña et al., 2019) |
| | *Bacillus megaterium* TRQ8, *B. cabrialesii* TE3T, *B. paralicheniformis* TRQ65 | Biochemical, growth and yield measurements | Optimal | (Rojas-Padilla et al., 2022) |

| | | | | |
|---|---|---|---|---|
| | Not indentified | Growth measurements and As adsoption | Optimal | (Xiao et al., 2020) |
| | *Acidovorax delafieldii* | Biochemical, growth and yield measurements | Optimal | (Cavite et al., 2021) |
| | *Bacillus sp.* | Biochemical, growth and yield measurements | Optimal | (Pan et al., 2023) |
| | *Enterobacter cancerogenus E50S2-4, Bacillus amyloliquefaciens E50S2-3, Klebsiella quasipneumoniae M50R2-3, and Bacillus velezensis M100S1-4* | Biochemical and growth measurements | Pesticide, Salt and drought stress | (Ma et al., 2023) |
| **Rice** | *Pantoea ananatis D1* | Biochemical and growth measurements | Salt stress | (Lu et al., 2021) |
| | *Bacillus thuringensis, Lysinibacillus sp. N5, Bacillus sp. N7* | Yield measurements | Ozone stress | (Autarmat et al., 2023) |
| | Alcaligenes sp. | Biochemical and growth measurements | Salt tolerance | (Fatima et al., 2020) |
| | *Bacillus pumilus TUAT1* | Growth measurements | Optimal | (Agake et al., 2022) |
| | *Bacillus pumilus JPVS11* | Biochemical and growth measurements | Salt stress | (Kumar et al., 2021) |
| | *Enterobacter cloacae CTWI-06* | Growth and yield measurements | Cr contaminated soil | (Pattnaik et al., 2020) |

Rojas Padilla, J., Chaparro Encinas, L.A., Robles Montoya, R.I., Santos Villalobos, S. de los, 2020. Growth promotion on wheat (Triticum turgidum L. subsp. durum) by co-inoculation of native

Bacillus strains isolated from the Yaqui Valley, Mexico. Nov. Sci. 12, 1–29.
Rozier, C., Hamzaoui, J., Lemoine, D., Czarnes, S., Legendre, L., 2017. Field-based assessment of the mechanism of maize yield enhancement by Azospirillum lipoferum CRT1. Sci. Rep. 7, 7416.
Safdarian, M., Askari, H., Nematzadeh, G., Adriano, S., 2020. Halophile plant growth-promoting rhizobacteria induce salt tolerance traits in wheat seedlings (Triticum aestivum L.). Pedosphere 30, 684–693.
Singh, S., Jagota, N., Kaur, H., Kaur, R., Kaur, G., Sandhu, S., Sharma, A., 2023. Deciphering behavioral changes in maize plants in a quest to identify species specific plant growth promoting rhizobacteria. Total Environ. Res. Themes 6, 100043.
Xiao, A.W., Li, Z., Li, W.C., Ye, Z.H., 2020. The effect of plant growth-promoting rhizobacteria (PGPR) on arsenic accumulation and the growth of rice plants (Oryza sativa L.). Chemosphere 242, 125136.
Zarei, T., Moradi, A., Kazemeini, S.A., Akhgar, A., Rahi, A.A., 2020. The role of ACC deaminase producing bacteria in improving sweet corn (Zea mays L. var saccharata) productivity under limited availability of irrigation water. Sci. Rep. 10, 20361.

#### Supplementary figures

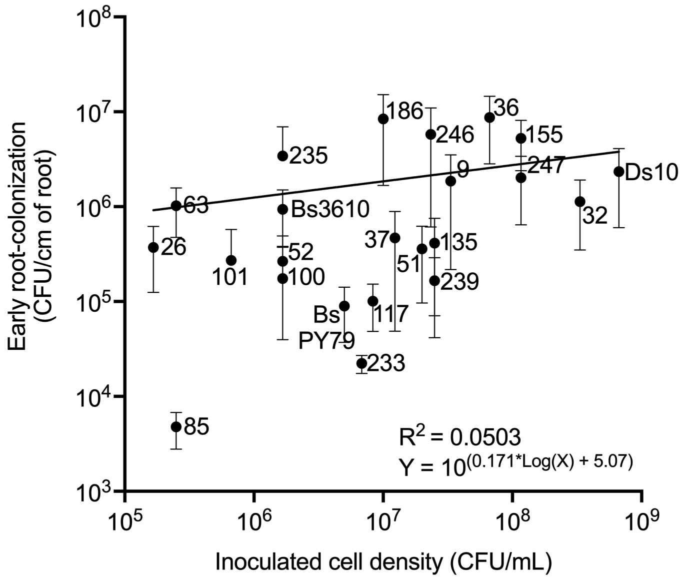


**Figure S1**. **Correlation between early root-colonization and inoculated CFUs.** Numbers beside each dot represent the code of NME strains. Log-log line derived from a non-linear regression analysis is shown in black. Error bars indicate 95% confidence intervals.

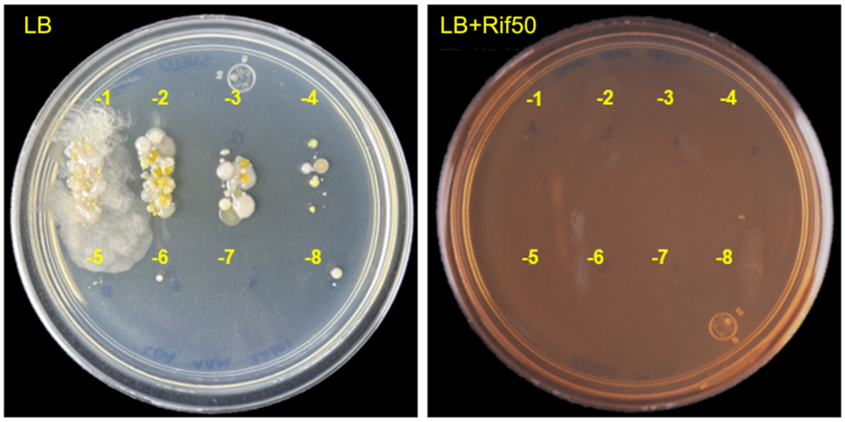


**Figure S2**. **Soil suspension spotted on LB and LB+Rif50.** A sample of 100 mg of soil was suspended in 500 µL of PBS. The suspension was vortexed for 40 s and let sit for 1 min. Then, 10 µL of 1:10 serial dilutions were spotted on LB and LB + 50 µg/mL Rifampicin. Pictures were taken after an overnight incubation at 30 ºC.

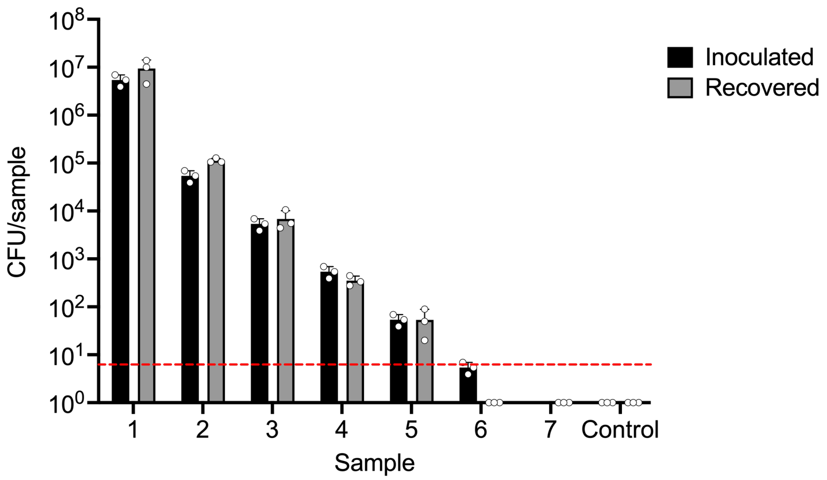


**Figure S3. CFU counts of inoculated soil samples.** Each sample was inoculated with descendant concentrations of NME_117 Rif$^r$. Sample 1 was inoculated with a non-diluted inoculum. Samples 2 to 7 were inoculated with concentrations from $10^{-2}$ to $10^{-7}$ of the same inoculum, respectively. CFUs were assessed from soil samples after an ON incubation at room temperature. The detection limit for this assay is shown with a red dotted line. Bars indicate mean ± SD.

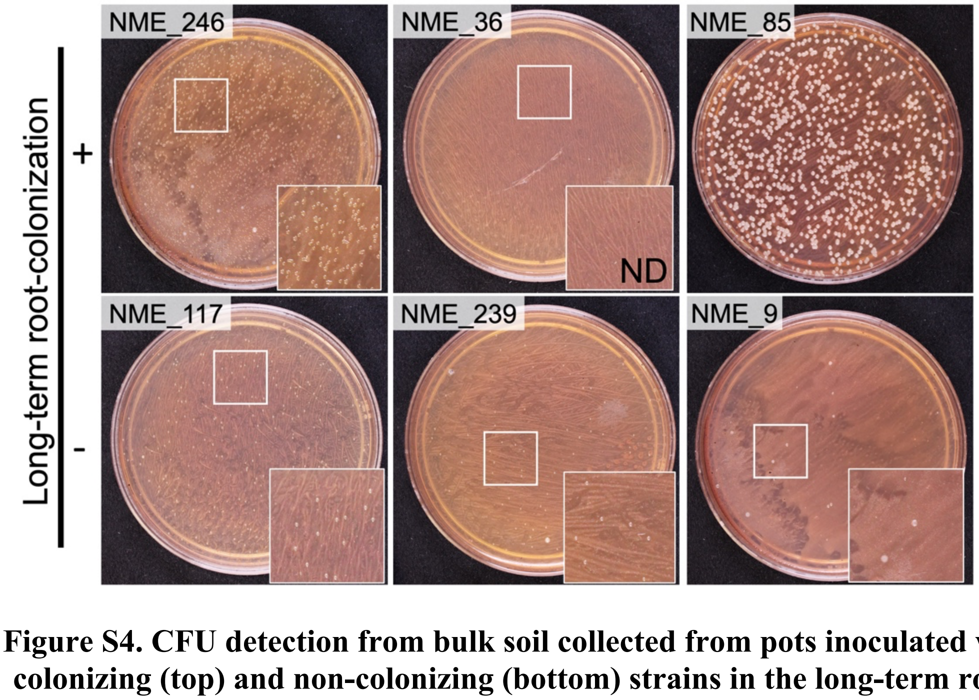


**Figure S4. CFU detection from bulk soil collected from pots inoculated with colonizing (top) and non-colonizing (bottom) strains in the long-term root-colonization assay.** Insets show zoomed-in sections of the corresponding Petri dish where colonial growth is evident. ND, colonies not detected.

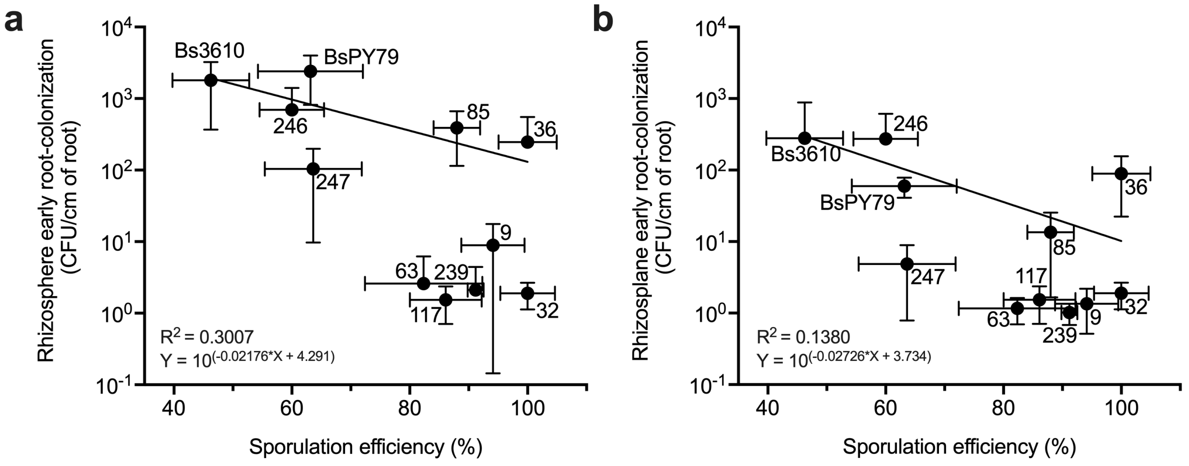


**Figure S5. Correlation between long-term rhizosphere (a) and rhizoplane (b) root-colonization and sporulation efficiency of each strain.** Numbers beside each dot represent the code of NME strains. Semilog line derived from a non-linear regression analysis is shown in black. Error bars indicate SD.

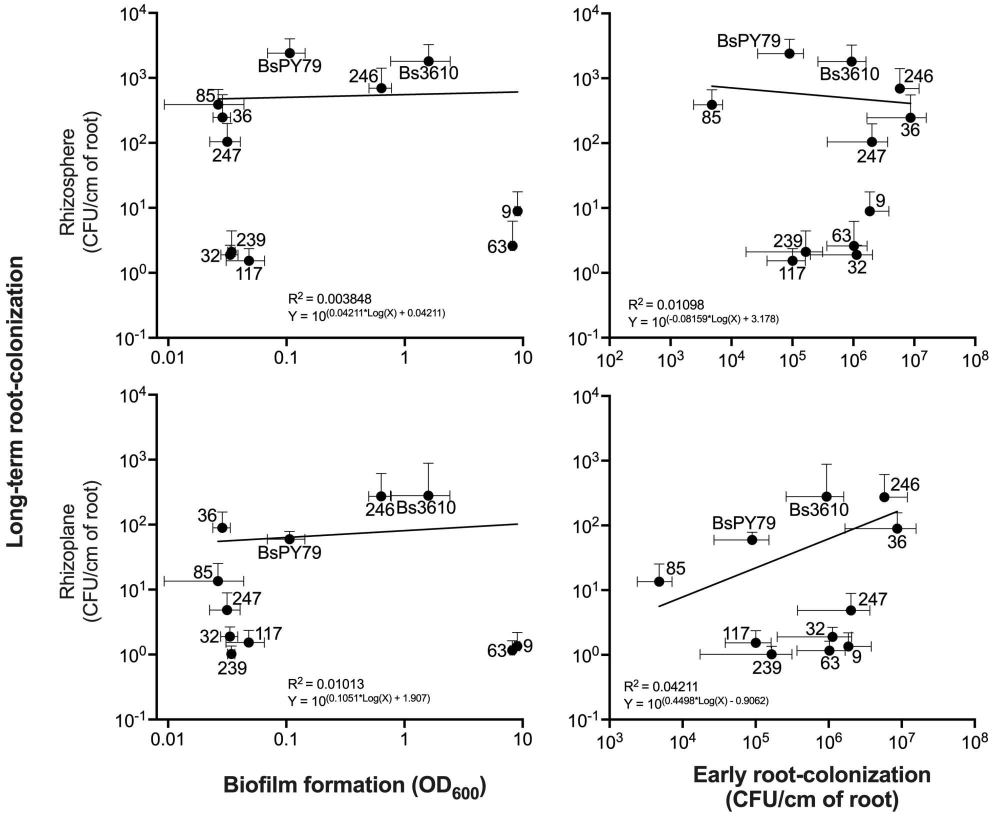


**Figure S6. Correlation between long-term root-colonization and biofilm formation (top and bottom left), and early root-colonization (top and bottom right).** Numbers beside each dot represent the code of NME strains. Log-log line derived from a non-linear regression analysis is shown in black. Error bars indicate SD.

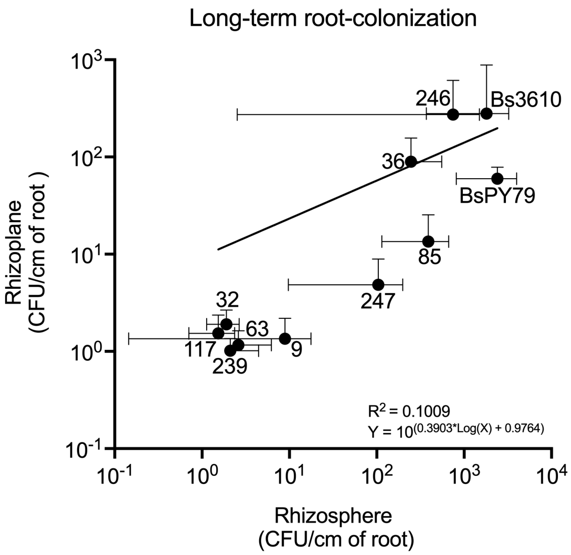


**Figure S7. Correlation between long-term rhizosphere and rhizoplane colonization.** Numbers beside each dot represent the code of NME strains. Log-log line derived from a non-linear regression analysis is shown in black. Error bars indicate SD.

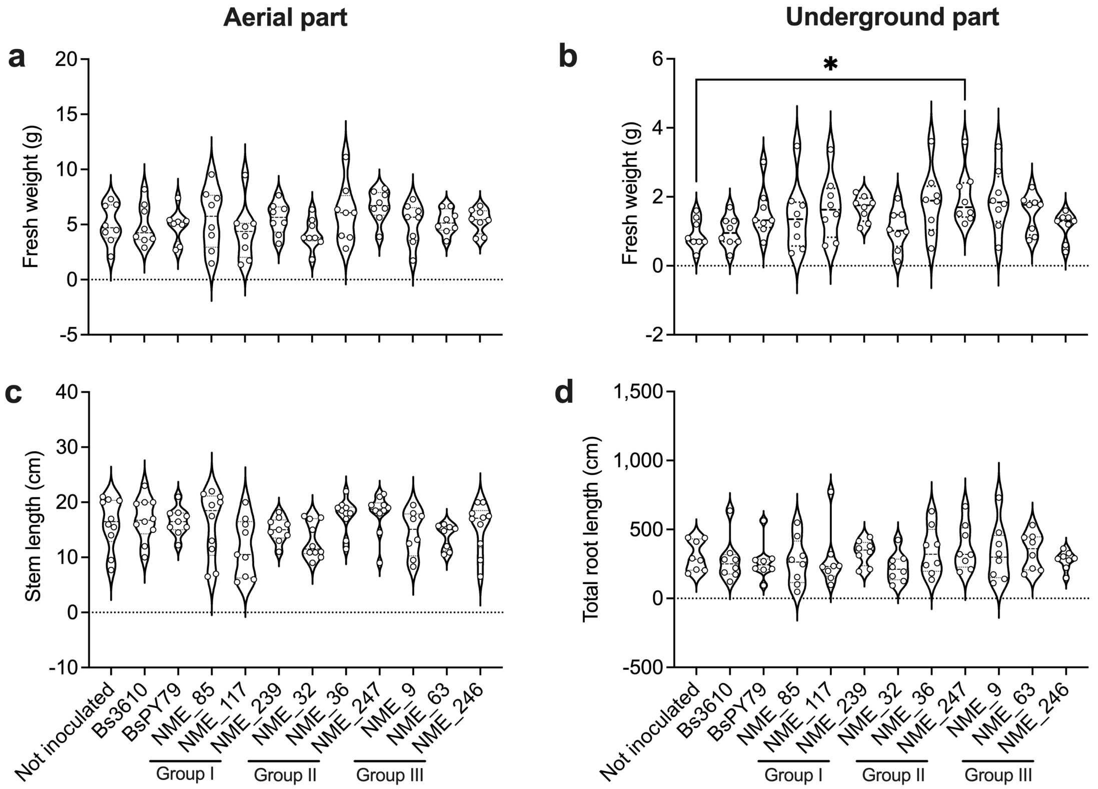


**Figure S8. Morphometric values of plants inoculated with seed-endophytic Bacilli from native maize landraces.** Differences between non-inoculated and inoculated plants were assessed using a one-way ANOVA and Dunnett test. *, $p<0.0332$.
